# Paternal kin discrimination by sons in male chimpanzees transitioning to adulthood

**DOI:** 10.1101/631887

**Authors:** Aaron A. Sandel, John C. Mitani, Kevin E. Langergraber

## Abstract

Although paternal investment explains the evolution of fatherhood from a functional perspective, its evolutionary origins are unclear. Here we investigate whether a building block for paternal investment, father-offspring discrimination, is present in our closest living relatives, chimpanzees. Adolescent and young adult males (12 - 21 years old) maintained proximity and groomed with their fathers more frequently than with other males given how often they associated. This discrimination did not likely increase the short-term inclusive fitness of fathers or sons because the absolute time they spent in proximity or grooming did not exceed the time spent in these activities by other dyads. Almost all grooming was done by sons rather than fathers, suggesting that sons are responsible for observed biases in father-son behavior. Father-offspring discrimination could partly be explained by young males socializing with males who were more likely to be their father based on their age at the time of the young male’s conception. Two other cues of paternity, the other male’s rank at the time of the young male’s infancy and the other male’s association frequency with the young male’s mother during the young male’s infancy/juvenility, failed to predict association-controlled proximity or grooming. Father-son biases persisted even after controlling for characteristics of males that predicted paternity probability, implicating other cues that we did not examine. These results suggest that an important factor for the evolution of highly investing fathers in humans, father-offspring discrimination, may have been present in simpler form in the last common ancestor they shared with chimpanzees.

## Introduction

In most human societies, fathers contribute to the health and fitness of their offspring. Through co-evolution with sexual-division of labor, this has led to the unusual combination of high fertility and low mortality responsible for our ecological dominance of the planet (1–6). Although human fatherhood has been well studied from this functional perspective, its evolution (sensu 7) remains puzzling. Phylogenetic inertia plays a strong role in the evolution of social and mating systems, with those of closely related species generally being very similar (8, 9). Yet in humans’ closest living relatives, chimpanzees and bonobos, females are highly promiscuous and mate with many adult males during each conception cycle (10–12). Like all complex adaptations, human fatherhood represents the end-product of a multi-step evolutionary sequence in which more elaborate forms succeeded one another (13). For fathers to invest heavily in their offspring, they must first evolve a capacity for kin discrimination, which refers to differences in behavioral responses individuals show toward kin compared to non-kin using conspecific cues correlated with kinship (14).

Pair bonds play a key role in father-offspring discrimination in all contemporary human societies. In these, fathers discriminate their offspring born by the woman with whom they had a pair-bond and more or less exclusive sexual access, and offspring discriminate their father as the man who had a long-term pair-bond with their mother (13, 15, 16). Although highly investing fathers seem to occur only in species with strong pair-bonds (e.g., titi monkeys: 17), a growing body of evidence shows that weaker father-offspring discrimination occurs in species where females mate promiscuously (18–24). This evidence includes two studies of chimpanzees suggesting that fathers show slight tendencies to play and groom with their own infants and juveniles (25, 26).

Several non-mutually exclusive cues can contribute to father/offspring discrimination in promiscuous species. Some cues are indirect and rely on the fact that even in the most promiscuous species paternity success is not entirely random but instead is correlated with certain male characteristics (14, 27, 28). For example, in the Ngogo community of chimpanzees, the probability that a male is the father of an offspring is predicted by his dominance rank at the time of conception and his long-term tendency to associate with the offspring’s mother (29). Biases in the social behavior between chimpanzee offspring and fathers could thus arise as a byproduct of offspring preferentially interacting with males who were high ranking and frequently associated with their mothers when offspring were young. Alternatively, a similar bias would result if fathers preferentially interact with offspring born to females with whom they frequently associated when they were high ranking. Baboons provide some of the strongest evidence for father-offspring discrimination in promiscuously mating mammals as females form special relationships with formerly high-ranking males who are likely to be the fathers of their offspring (18, 21, 30, 31).

Age is another indirect cue for paternal kin recognition in promiscuous species. In several female-philopatric primate species, adult females similar in age tend to socialize. These individuals are likely to be paternal sisters because only a small number of males reproduce at any given time (32–34). However, unlike paternal siblings, fathers and offspring are by necessity very different rather than similar in age, as paternity success typically has an inverted-U relationship with age, increasing from sexual maturity, peaking in prime adulthood, and declining thereafter (35). Whether animals use differences in age as a cue of a father-offspring relationship is unknown.

Father-offspring discrimination may also involve direct cues (sensu 14, 27) as occurs in phenotype matching, where individuals discriminate among conspecifics based on their perceived phenotypic and thus genetic similarity to themselves or known kin. Evidence for phenotype matching has been demonstrated experimentally (36, 37), but more often is invoked when other indirect cues have been excluded, especially in field studies of animals in the wild (28, 38).

In this paper, we investigate father-offspring discrimination and its underlying cues in wild chimpanzees. Unlike previous research on kin discrimination in promiscuous species, which focused on behavior between fathers and sexually immature offspring (i.e., infants and juveniles), we examine behavior between fathers and sexually mature offspring (i.e., adolescents and young adults). We focus on male offspring because they remain in their natal community, and thus unlike females who disperse, have opportunities to socialize with their fathers well past sexual maturity. Opportunities for sexually mature males to interact with their fathers are especially likely in our particular chimpanzee study group at Ngogo in Kibale National Park, Uganda, where minimal human disturbance and an abundant food supply support an unusually large community (N = 193 individuals), whose individuals survive at high rates and live a long time (39).

During 12 months we recorded the amount of time 18 adolescent and young adult males (‘young males,’ age range: 12-21 years) spent with one another and with 36 other younger adolescent and older adult males (8 - 53 years, including 11 fathers) in: (1) association in the same party; (2) spatial proximity of ≤ 5 m; and (3) grooming. While proximity and grooming are widely considered to be meaningful measures of affiliative social behavior in non-human primates, their analysis is complicated in fission-fusion species like chimpanzees, where dyads vary in their opportunities to perform these behaviors based on how often they associate in temporary subgroups or ‘parties’ (40). Association, in turn, reflects social preferences, but also additional social, ecological, and other constraints (41). We compare levels of proximity and grooming in father-son and other types of dyads with and without controls for frequency of association, interpreting the former as an indication of discrimination and the latter as an indication of the potential positive impacts of discrimination on inclusive fitness (34, 42, 43).

## Results

### Kin discrimination

We found evidence for father-son discrimination: when associating in the same party, young males were in proximity and groomed with their fathers more frequently than with ‘other’ males (Table 1). The ‘other’ category includes unrelated males and also paternal brothers, uncles, cousins, and additional, more distant kin that, unlike maternal brothers, do not preferentially socialize in male chimpanzees (44, 45; Figure 1). Consistent with previous research that employed a broader age range of dyads, i.e., not limited to dyads containing at least one young male (44, 45), young males were in proximity with their maternal brothers more often than with other males when in association (Figure 1). In contrast to prior studies, young males did not groom with their maternal brothers more frequently when associating than other males did when together. Young males actually groomed with their fathers as often as with their maternal brothers when associating, although they were less often in proximity (Table 1).

**Table 1.**
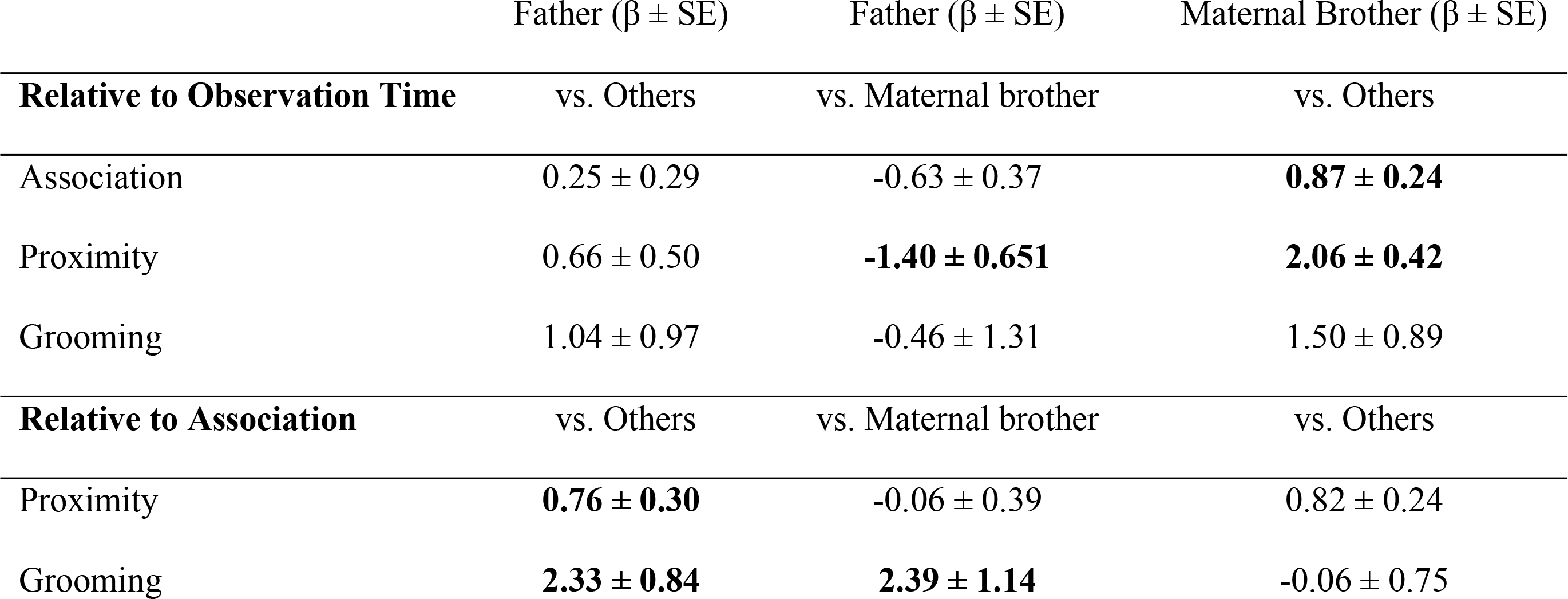
Kin discrimination in social behavior. Coefficients for the effect of fathers and maternal brothers on dyadic behavior compared to ‘other’ males (including paternal half-bothers and distantly or unrelated males). Informative predictors are represented in bold typeface. Visual representation, including individual data points, is represented in Figure 1.

| | Father ($\beta \pm SE$) | Father ($\beta \pm SE$) | Maternal Brother ($\beta \pm SE$) |
| --- | --- | --- | --- |
| **Relative to Observation Time** | vs. Others | vs. Maternal brother | vs. Others |
| Association | 0.25 ± 0.29 | -0.63 ± 0.37 | **0.87 ± 0.24** |
| Proximity | 0.66 ± 0.50 | **-1.40 ± 0.651** | **2.06 ± 0.42** |
| Grooming | 1.04 ± 0.97 | -0.46 ± 1.31 | 1.50 ± 0.89 |
| **Relative to Association** | vs. Others | vs. Maternal brother | vs. Others |
| Proximity | **0.76 ± 0.30** | -0.06 ± 0.39 | 0.82 ± 0.24 |
| Grooming | **2.33 ± 0.84** | **2.39 ± 1.14** | -0.06 ± 0.75 |

**Fig 1.**
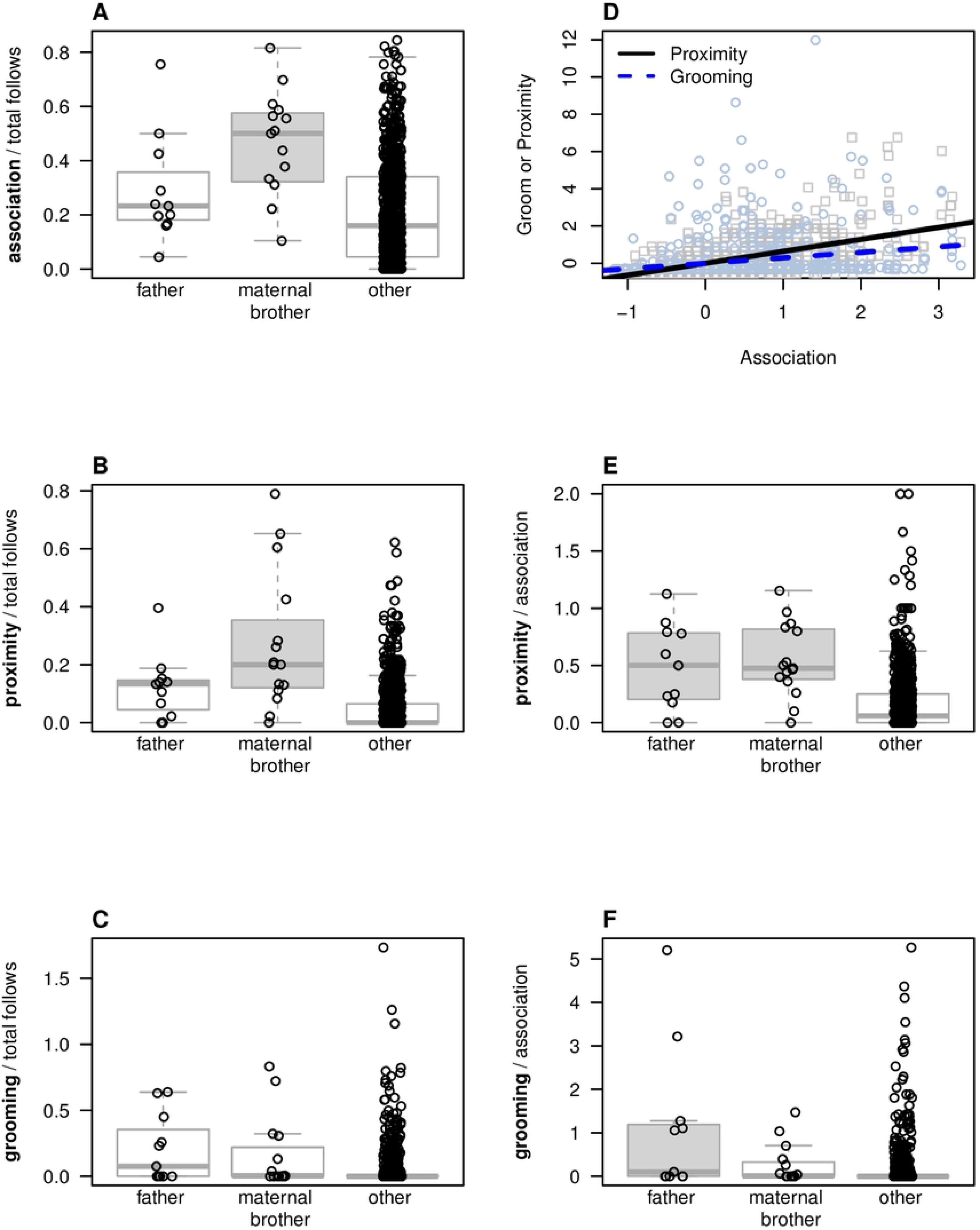
Social behavior patterns based on relatedness. (a) association, (b) proximity, and (c) relative to total observation time, and (e) proximity and (f) grooming relative to total association time; (d) correlation between association and proximity and grooming. Shaded boxes indicate kin categories that had positive coefficients that did not overlap with zero; statistical tests were done using multiple regression of relatedness.

### Potential inclusive fitness effects of kin discrimination

Although young males were more likely to be in proximity and groom with their fathers when associating, it is unlikely that this discrimination increased the inclusive fitness of either member of the dyad (at least in the short-term – see discussion), as absolute levels of proximity and grooming among father-son dyads were comparable to those between other dyads (Figure 1). Nor did young males associate in parties more frequently with their fathers than with other males. While it is possible that these null results represent false negatives due to lower power from our small sample (N = 11 father-son pairs), this explanation is unlikely given that we also found that young males preferentially associated and maintained proximity (but did not groom) with their maternal brothers (N = 15 pairs) in a similarly sized sample.

### Cues underlying father-son discrimination

We now turn to analyses designed to identify the cues underlying the tendency for males to be in proximity and groom with their fathers when associating with them. Our first step was to determine which member of the pair was responsible for father-son discrimination. Although we did not collect data that would allow us to address this question for proximity, it was clear that sons were responsible for the tendency of father-son dyads to groom when associating in parties. Among 11 father-son dyads, 6 sons groomed their fathers, and the average number of seconds (± SD) sons groomed their fathers per son focal hour was 12.7 ± 15.4. In contrast, only one father ever groomed his son, and the average number of seconds fathers groomed their sons per son focal hour was 0.2 ± 0.7.

Under the assumption that sons rather than fathers are responsible for the tendency of these dyads to be in proximity and groom when associating, we next investigated how these kin biases could arise if young males socialized with other males based on characteristics statistically associated with paternity probability. In our first set of models (‘all ages’ models), which employed the same age range of potential partners of young males as above, we examined how the 18 young males’ association-controlled proximity and grooming with 53 other males aged 8-53 in 2014-2015 were independently predicted by the other males’: (1) rank when the young male was an infant; (2) probability of being the younger male’s father based on his age at the time of the young male’s conception (i.e., male age-specific fertility, estimated using 15 years and N = 105 paternities from long-term Ngogo data (29, 44–46)); and (3) whether the other male actually was the young male’s father. Our reasoning for including this third predictor variable was that if father identity explains additional variation in young adult/potential father social behavior even after accounting for indirect cues for paternity probability, this would suggest that additional cues not included in our model may be involved in father/son discrimination. In a second set of models (‘potential sire age only’), we examined how the 18 young males’ association-controlled proximity and grooming were independently predicted by the same predictors as in the models above, plus an additional predictor variable: (4) the other male’s association frequency with the young male’s mother when the young male was an infant or juvenile (~11 years previously in 2003-2004, the only period for which such archival data were available). In this second set of models we only included 21 older males aged 22 - 53 years in 2014-2015 as potential social partners of the 18 young males. Only males of this age were ≥ 10 years old and reproductively active in 2003-2004 and could thus increase their chance of siring the latter by associating with his mother during this time (29).

Young males did not maintain proximity or groom when in association with males who were high ranking at the time of the young male’s infancy (Figure 2). Nor did young males have high levels of association-controlled proximity or grooming with older males who had frequently associated with the young male’s mother while he was an infant or juvenile. In contrast, there was some evidence that father-son discrimination could be partly explained by the tendency for young males to socialize with males who had high age-specific fertility at the time of the young males’ conception. This variable predicted association-controlled grooming in the ‘all ages’ model, but not in the ‘potential sire age only’ model, nor did it predict association-controlled proximity in either model (Figure 2). In both models, young males had higher association-controlled proximity and grooming with their fathers than with other males even after controlling for characteristics of males that predicted paternity.

**Fig 2.**
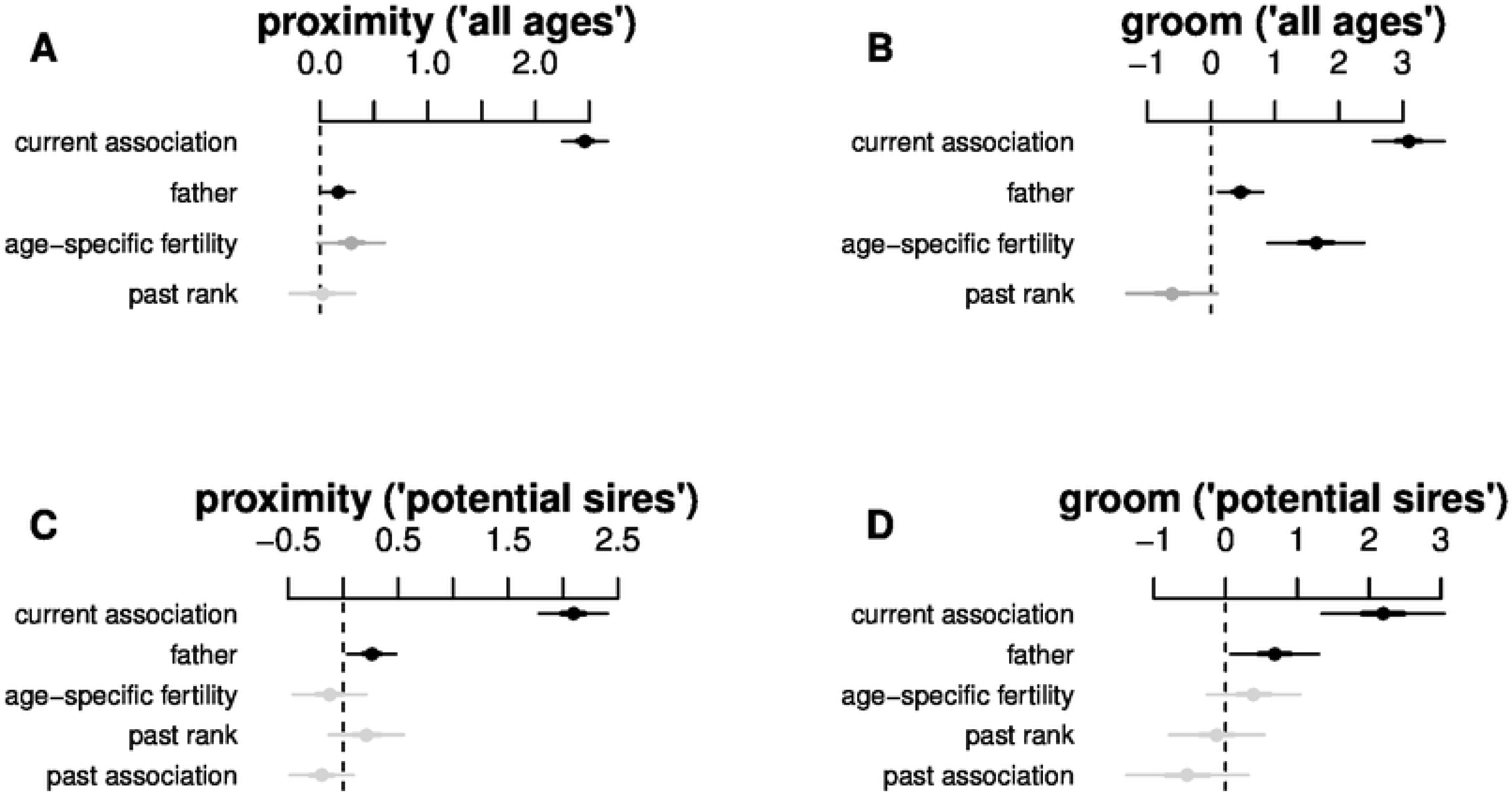
Coefficients of potential cues of father-son discrimination from four multivariate models. Thick lines represent ±1 SD and thin lines for ±2 SD. Informative predictors are represented in black.

## Discussion

We found evidence of father-son discrimination in chimpanzees, as adolescent and young adult males (‘young males’) and their fathers were in close spatial proximity and groomed more often when associating than did other dyads. While these results add to a growing body of studies revealing father-offspring discrimination in promiscuously mating, group-living species, they differ from most previous work in that they involved discrimination by offspring rather than by fathers. This finding may seem surprising given Hamilton’s (47) ‘fundamental asymmetry,’ which suggests that because the benefits of receiving help should be higher for offspring and the costs of giving help should be lower for parents, nepotism should be directed from parents to offspring rather than the reverse. Hamilton’s asymmetry, however, is based on the assumption that actors perform behaviors for related recipients’ because these behaviors increase recipients’ direct fitness, and thus the actor’s indirect fitness. Although our results suggest that young males discriminated their fathers, it is unlikely that this discrimination actually resulted in direct fitness gains for fathers (and thus indirect fitness gains for young males), as the absolute amount of time that young males were in proximity with and groomed their fathers was low, and not significantly different from other males. Thus, father-son discrimination may occur because it results in longer-term, direct fitness gains for sons.

How might young males increase their direct fitness by preferentially maintaining proximity with and grooming their fathers when associating with them in parties? Part of the tendency for males to groom their fathers when in association could be explained by young males performing this behavior with males whose age difference from themselves corresponded with the probability of fatherhood. At Ngogo, over 50% of offspring were sired by males age 18-26, and the mean ages of fathers of young males in the current study was 40.4 years (range = 33.7 – 45.7 years). Socializing with these older, past-prime aged males may help young males by facilitating their entry into the social network of adult males (11). During infancy and juvenility, males are in near constant contact with their mothers, who serve as their primary grooming partner and source of support (48–51). This changes drastically during adolescence, when males begin to travel independently of their mothers and socialize with adult males. As they make this transition, adolescents receive increased aggression from adult males (50) and continue to remain at the periphery of parties, sometimes even after reaching adulthood (52). As a consequence, adolescent males appear to be “less relaxed” and more “tense and inhibited” when they are around adult males (Pusey 1990: 228). While old males are socially integrated into the network of adult males, they are generally lower ranking, less aggressive, and more tolerant of young males than are prime-aged males (53–57). This interpretation is consistent with research in other species suggesting direct fitness benefits may sometimes play a larger role than indirect fitness benefits in the evolution of kin discrimination (e.g., 58, 59, 60). For example, in many group-living primates, individuals preferentially cooperate with individuals who are similar in age, as cooperating with individuals who have similar needs, access to resources, and abilities to exchange them results in the highest direct fitness (45, 61, 62). When patterns of male reproductive skew result in age-mates being paternal siblings, individuals gain additional indirect fitness benefits by socializing with age-mates as a byproduct of striving to maximize their direct fitness (32).

Not all son-father discrimination could be explained by young males preferentially socializing with old males, as association-controlled proximity and grooming was elevated in young males and fathers even when controlling for the age difference between the pair. While similar findings in previous studies of kin discrimination have been tentatively interpreted as evidence for phenotype matching (33, 63), we must acknowledge the limitations of our cue analyses. Our sample of father-son pairs was small (N = 11 pairs), limiting the power of our multivariate analyses and the number of predictor variables we could include. For example, we found that association-controlled proximity and grooming patterns of young males were not predicted by the other males’ association frequency with the young males’ mother when the young male was an infant or juvenile, but this measure of mother–potential father behavior was based on only one year and one type of behavioral data. Observations across their entire period of development and incorporating additional measures of social behavior between potential fathers and the mothers of young males beyond association, such as proximity or grooming, might yield additional insights into the cues that young males use to discriminate their fathers.

Irrespective of the underlying cues on which father-son discrimination is based, our finding that father-son discrimination occurs in chimpanzees goes some way to bridge the gap between the social and mating systems of humans and their closest living relatives. These results provide support for Chapais’ (13: 199) suggestion that if human fatherhood does have phylogenetic building blocks in a chimpanzee-like society, “enduring father-son bonds might have been initiated and maintained by the sons themselves, hence independently of and prior to the evolution of active forms of paternal care.” As Chapais notes, complex adaptations like human fatherhood represent the end product of a multistep evolutionary sequence where progressively more elaborate versions succeeded one another, and the present-day adaptive function of a trait may be a poor guide for its earlier evolutionary origins.

## Methods

### Behavioral observations

A.A.S. conducted behavioral observations of adolescent and young adult males during focal sampling sessions lasting one hour (mean ± SD hours of observation per subject = 43 ± 3.2 hours, N = 18 males) from 24 August 2014 to 30 August 2015, collecting data on association, proximity, and grooming. Males who encountered one another for any length of time during hour-long following episodes were scored as in association with the focal subject. Individuals in proximity (≤ 5 meters) to the focal subject were recorded during instantaneous point samples made at 10-minute intervals. The amount of grooming given and received by focals was recorded to the nearest second. Research was reviewed by the University Committee on Use and Care of Animals at the University of Michigan, and was exempt because animal use was limited to non-intrusive field observations.

To assess whether social relationships of adolescent and young adult males in 2014-2015 were related to social relationships they had in the past as infants and juveniles, we examined adolescent and adult female and male party associations during 2003 and 2004, the one period for which these data were available and collected by K.E.L (29, 64). Instantaneous point samples were made at half hour intervals to record adolescent and adult males in association with focal adult female subjects. Because infant and juvenile chimpanzees are in near constant contact with their mothers (49) maternal party associations reflect those of their infant and juvenile sons. We evaluated associations between mothers and potential fathers using the half-weight index (65) in SOCPROG (66).

To assess the dominance status of older males, we used observations of pant grunts, a formal signal of submission directed up the hierarchy and given by low-ranking chimpanzees to higher-ranking individuals (67). Pant grunts exchanged between males were recorded by J.C.M. between 1995 and 2004. To determine the past dominance rank of older males, we combined pant grunts exchanged between males into a single giver-receiver matrix. Males were given ordinal ranks (e.g. the alpha male had a rank of 1, the beta had a rank of 2). Since ordinal ranks have different meaning depending on the total number of males in a given year, we controlled for the number of males by subtracting the rank from the total number of males in the hierarchy, and divided this by the total number of males minus one. Thus the highest-ranking male had a rank of 1 and the lowest-ranking male had a rank of 0 (68).

### Kinship

Genetic relationships between all of our subjects are known based on prior behavioral observations and genetic analyses of autosomal, X-chromosomal, and Y-chromosomal microsatellite loci, and of mitochondrial DNA (29, 44, 46, 64). Eleven adolescent and young adult males had fathers who were alive at the time of this study; fathers included six males. Fifteen adolescent and young adult males had maternal brothers who were adolescent or adults during the time of this study.

### Male age-specific fertility

For every offspring born with a known father between 1 Jan 2003 and 12 Feb 2018, we determined the age of the father at the time of conception, rounded to the nearest year. We limited the data to infants born in or after 2003 because by this time, nearly all infants born each year were genotyped. Paternities were not available for 52 of 159 infants born during this period: 23 infants died within 2 months of birth, and the genotypes of 9 infants born between 2017 and 2018 have not yet been determined. To determine age-specific fertility, we calculated the percentage of offspring sired at each male age. We then calculated a 5-year running average. For example, for 20-year-old males, this included the average percentage of offspring sired by 18-year-old males, 19-year-old males, 20-year-old males, 21-year-old males, and 22-year-old males. The running average became the male age-specific fertility value. We assigned this value to each dyad based on the older male’s age at the time of the younger male’s conception.

### Statistical Analyses

To assess social behavior between male chimpanzees, we analyzed associations, proximity, and grooming interactions separately. While some researchers combine different affiliative behaviors into a single index (42, 69), each behavior may reflect different aspects of social behavior (70, 71) and combining them may not always be appropriate (72). Association and proximity were kept as counts. Grooming was measured as a continuous variable as the duration of time dyads spent receiving and giving grooming summarized across the entire year.

To assess the effects of kinship and examine whether a bias to socialize with fathers existed, we conducted three generalized linear mixed models (GLMMs), with association, proximity, and grooming between pairs of males as the outcome variables. Fixed effects were the dyads’ kin relationship (i.e., father-son, maternal brothers, or ‘other’). The ‘other’ category was set as the reference class. First, we constructed two models for proximity and grooming in which we excluded dyads that never associated and added the log number of times each pair was in association as a fixed effect to control for variation in opportunities to interact. Second, we assessed absolute levels of association, proximity, and grooming for which we added the log number of hour-long following episodes on the focal subject as a fixed effect to control for variation in observation time. Third, to assess kin discrimination mechanisms, we constructed four GLMMs, with grooming or proximity between adolescent and young adult males and other males in 2014-2015 as the outcome variables. Predictors are outlined in the main text above.

All fixed effects were centered and z-transformed to increase interpretation of relative variable importance (73). In all models, the identities of subjects and partners were included as random effects. We set a negative binomial error distribution using the ‘glmmADMB’ package (74) in R (75). We report coefficients and standard errors for each kinship category as an indicator of variable importance. We considered a predictor to be informative if its coefficient minus twice its standard error did not include zero.

## Acknowledgements

Fieldwork was sponsored by the Uganda Wildlife Authority, the Uganda National Council for Science and Technology, and the Makerere University Biological Field Station. Nathan Chesterman provided assistance in the field. For additional support, we are grateful to David Watts, Sam Angedakin, Alfred Tumusiime, Ambrose Twineomujuni, Godfrey Mbabazi, Laurence Ndangizi, Rachna Reddy, and the late Jerry Lwanga. For statistical advice, we thank László Garamszegi and the Center for Statistical Consulting and Research at the University of Michigan, especially Kerby Shedden. For helpful feedback during the development of this project and comments on earlier versions of the manuscript, we thank Jacinta Beehner, Thore Bergman, Anne Pusey, Brent Pav, Rachna Reddy, Sam Patterson, Bethany Hansen, Barb Smuts, Kiku Adatto, Michael Sandel, Joan Silk, and additional friends, family, and colleagues of A.A.S.

## Supporting information

**S1 File. Current social behavior**. Dyadic data for adolescent and young adult males (12 – 21 years) with all other adolescent or adult males (≥8 years).

**S2 File. Behavioral mechanisms (‘all ages’)**. Dyadic data for adolescent and young adult males (12 – 21 years) with other adolescent or adult males (≥9 years).

**S3 File. Behavioral mechanisms (‘potential sires only’)**. Dyadic data for adolescent and young adult males (12 – 21 years) with older adult males (≥22 years).

